# Self-regulation of attention in children in a virtual classroom environment: a feasibility study

**DOI:** 10.1101/2023.08.02.551583

**Authors:** Carole Guedj, Rémi Tyrand, Emmanuel Badier, Lou Planchamp, Madison Stringer, Myriam Ophelia Zimmermann, Victor Férat, Russia Ha-Vinh Leuchter, Frédéric Grouiller

## Abstract

Attention is a crucial cognitive function that enables us to selectively focus on relevant information from the surrounding world to achieve our goals. When this sustained ability to direct attention is impaired, individuals face significant challenges in everyday life. This is the case for children with Attention Deficit Hyperactivity Disorder (ADHD), a complex neurodevelopmental disorder characterized by impulsive and inattentive behavior. While psychostimulant medications are currently the most effective treatment for ADHD, they often come with unwanted side effects, and sustaining the benefits can be difficult for many children. Therefore, it is imperative to explore non-pharmacological treatments that offer longer-lasting outcomes. Here, we proposed a groundbreaking protocol that combines electroencephalography-based neurofeedback (EEG-NFB) with virtual reality (VR) as an innovative approach to treating attention deficits. By integrating a virtual classroom environment, we aimed to enhance the transferability of attentional control skills while simultaneously increasing motivation and interest among children. The present study demonstrated the feasibility of this approach through an initial assessment involving a small group of healthy children, showcasing its potential for future evaluation in children diagnosed with ADHD. Encouragingly, the preliminary findings indicated high engagement rates and positive feedback from the children participating in the study. Additionally, the pre-and post-protocol assessments using EEG and fMRI recordings appeared to converge towards an improvement in attentional function. Although further validation is required to establish the efficacy of the proposed protocol, it represents a significant advancement in the field of neurofeedback therapy for ADHD. The integration of EEG-NFB and VR presents a novel avenue for enhancing attentional control and addressing behavioral challenges in children with ADHD.

## INTRODUCTION

Paying attention is not always easy, and lapses can occur even with our best efforts. Some of these failures are minor, like forgetting someone’s name or missing an object in our visual field. But others can have disastrous consequences, like distracted driving that leads to car accidents. While attention lapses are common, some people may have an amplified pattern of these failures, such as those with attention deficit hyperactivity disorder (ADHD). This disorder affects 5 to 7% of school-aged children (Thomas et al., 2015), and can lead to academic failure, anxiety, depression, behavioral disorders, relational issues, and drug addiction (Mochrie et al., 2020). Research has revealed that people with ADHD have reduced functional connectivity between frontal and parietal regions (Cocchi et al., 2012; Oldehinkel et al., 2016), and their brain activity shows increased levels of slow theta (4-8 Hz) activity and reduced alpha (8-12 Hz) and beta (13-30 Hz) activity (Rostami et al., 2022).

Conventional treatments for ADHD rely on stimulant medications such as methylphenidate (Ritalin®)(Sharma and Couture, 2014). These drugs work by increasing levels of norepinephrine and dopamine in the brain, which are considered the neurobiological cause of ADHD. However, 44% of children do not achieve remission (Swanson et al., 2011), many suffer from undesirable side effects (Storebø et al., 2018), and most have difficulty maintaining benefits beyond two years of treatment (Jensen et al., 2007). In addition, adherence to therapy, which is a crucial factor in the success of any treatment, is often a difficulty for children with ADHD entering adolescence (Kamimura-Nishimura et al., 2019).

An alternative approach called electroencephalography-based neurofeedback (EEG-NFB) has shown promise as it acts on healing and prevention, limits side effects, and potentially achieves lasting clinical benefits. With EEG-NFB, patients learn to regulate their brain electrical activity through real-time feedback extracted from their EEG signals. The ratio between theta and beta rhythms of the EEG has shown encouraging results in teaching children with ADHD to regulate their brain activity and reduce symptoms in the long term (Arns et al., 2009; Van Doren et al., 2017). Preliminary studies show that effects are maintained over follow-up periods of 6 and 24 months, with a tendency towards a greater decrease in hyperactivity and impulsivity symptoms after 24 months (Arns and Kenemans, 2014). One of the significant limitations of current approaches is that they require repetitive training sessions before stable and positive effects are achieved (Bink et al., 2015). In practice, this corresponds to 30-40 sessions involving long procedures such as the placement of numerous electrodes and the testing of their signal, followed by repetitive computer tasks that are difficult to maintain even for adults. The typical tasks of NFB protocols have nothing to do with how we sustain attention in real-life situations and with the challenges that children with ADHD face in their daily routines, especially in the classroom, which probably limits the transferability of what is learned during the intervention.

To overcome these limitations, we propose an innovative EEG-NFB protocol that leverages a virtual classroom environment. This environment has been specifically designed to enhance the transferability of attentional control skills beyond the training sessions and to challenge children through a distraction-modulated setting. By incorporating virtual reality, we aim to foster increased motivation and interest among the children, ultimately reducing the overall number of required sessions. In this report, we have demonstrated the feasibility of this groundbreaking protocol on a small group of healthy children aged 6 to 11 years. This study presents a significant step forward in neurofeedback treatment by validating an innovative and engaging protocol that holds promising potential for future testing in children with ADHD. This research opens up new horizons for the field, highlighting the effectiveness of a playful and immersive approach to neurofeedback therapy.

## METHODS

### 1. General procedure

The experimental protocol included a total of 12 sessions: 8 sessions of EEG-NFB based on theta/beta ratio (TBR) and performed in an immersive virtual classroom environment, 2 sessions of neuropsychological assessments (pre-and post-training) and 2 sessions of EEG-fMRI (pre-and post-training). The details of each session are summarized in Figure 1. Sessions were distributed in a maximum duration of 12 weeks with an optimal rate of one session of one hour per week.

**Figure 1.**
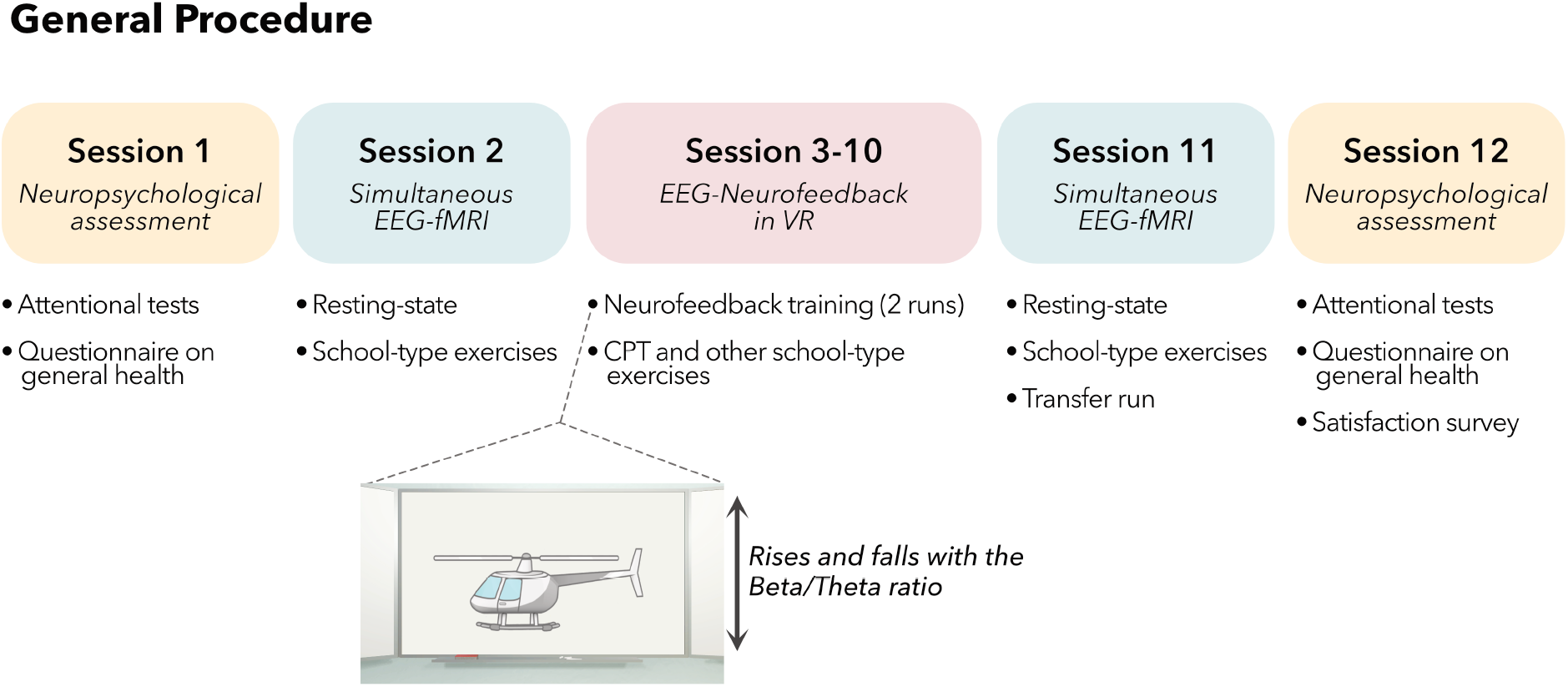

### 2. Participants

The study included a total of six children, with an average age of 9.46 years (SD: 1.23) and one female participant. However, two children were excluded from the final analysis due to technical issues during the neurofeedback training sessions. Additionally, another child was unable to complete the pre-and post-fMRI sessions due to artifacts caused by a dental appliance, but was still included in all other parts of the study.

To sum, a total of four children took part in EEG-NFB training sessions performed in a VR cave (sessions 3 to 8), and three of them also underwent EEG-fMRI scans before and after these sessions (sessions 2 and 11, respectively). For the EEG-NFB training data, this preliminary study analyzed the children’s performance during the Continuous Performance Task (CPT), as well as changes in the TBR. For the pre-and post-imaging sessions, we only analyzed the resting-state runs to evaluate data quality, and the transfer run to investigate the effect of attentional training on EEG power and brain activation.

The study was conducted in accordance with the Declaration of Helsinki and approved by the Research Ethics Committee of the Geneva University Hospital (CCER 2020-02642).

### 3. Material

#### a. Virtual classroom

##### i. General aspect

The virtual classroom environment was designed to closely mimic the daily reality of school children. The child was seated at a real school desk and immersed in the virtual environment, which included a classroom-like setting, such as a whiteboard, clock, and alphabet book on the wall (as depicted in Figure 2), as well as other students, a teacher, and a multitude of audio and visual distractions.

**Figure 2.**
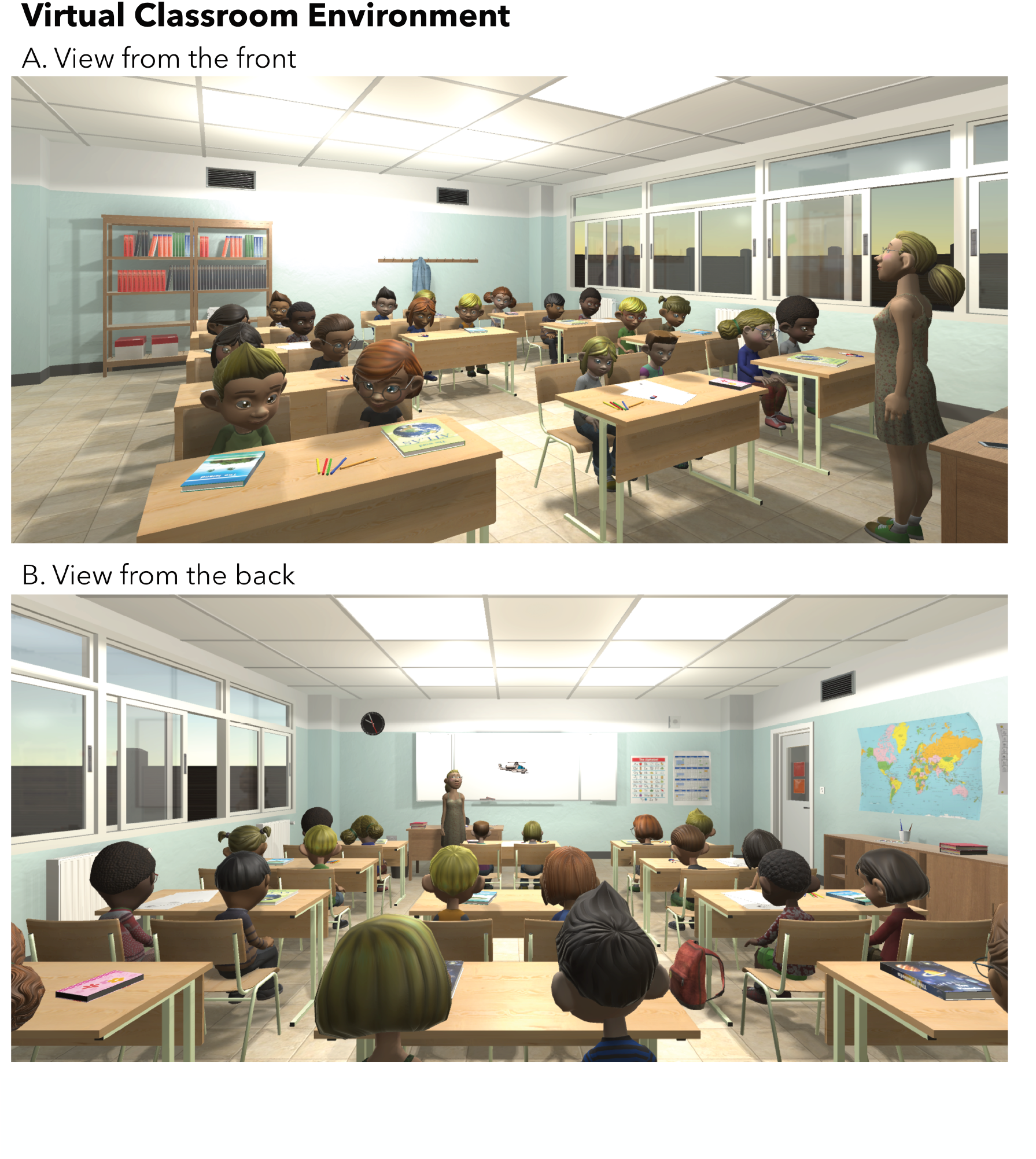

##### ii. Detailed equipment

The virtual classroom has been set up in the Brain Behavioral Laboratory Immersive-System (BBL-IS). This virtual reality (VR) cave system features four screens presenting seamless and perspective-coherent 3D images. The projection screens are made of four semi-transparent acrylic coated screens (Left, Front, Right, Ground). Their dimensions are 2.4m x 2.4m, except for the front screen which is wider (2.8m). These large screens enable the VR environment to be video-projected all around the user, enhancing the immersive experience. To do so, four Barco F70-4K8 video projectors were used with a resolution of 2560*1600 pixels and a refresh rate of 120Hz. Each video projector is located behind its corresponding screen to prevent the user from obstructing the projections. The VR cave also includes a motion tracking system composed of ten Vicon Bonita infrared cameras. Vicon Tracker software is used to track motion of the reflective markers positioned on the head of children. Vicon Tracker has a built-in Virtual Reality Peripheral Network server (https://vrpn.github.io/) which streams head positions to the VR simulation at a frequency of 240Hz. To deliver 3D spatialized sound, we used a 5.1 sound system (Denon AVR-1312, and KEF KHT2005.2 speakers).

The setup includes two computers. The first computer, featuring dual Intel Xeon E5645 CPUs, an Nvidia Quadro K5200 GPU, and 18 GB DDR3 RAM, serves for managing all recording systems such as motion tracking, EEG recording and real-time processing. The second computer, featuring dual Intel Xeon Gold 6144 CPUs, dual Nvidia Quadro RTX 6000 GPUs, and 128 GB DDR4 RAM, is dedicated to running the VR simulation. A MMBT-S Trigger Interface Box (Neurospec AG, Switzerland) is connected to this latter computer to synchronize the VR simulation with the EEG system. The code used was written in C-sharp and is available (https://github.com/ebadier/NeurospecTriggerBox-Unity). Finally, a 4- button Response Box (Current Designs Inc, PA, USA) was used to allow children to answer questions during tasks performed in VR.

##### iii. VR simulation

The VR simulation has been developed using the Unity game engine. We used the assets described in Table 1 from the Unity Asset Store.

##### iv. Script animation

The virtual scene was designed to closely mimic real-life classroom conditions and challenge the children’s attention levels in a progressive manner by incorporating distractions. The distractions were randomly introduced during specific time blocks of the VR simulation (every 30 seconds). Each task began with a 30-second distraction-free block. The frequency of distractions was proportional to the child’s level of concentration, as determined by the normalized TBR. The less focused the child was, the fewer distractions occurred, with a minimum of one distraction during distraction periods and a maximum of one distraction every 3 seconds if the child was fully focused. When not in distraction periods, the virtual characters displayed slow motion animations, but remained quiet. The distractions were randomly chosen from those described in Table 2 and only one distraction could occur at a time during distraction periods.

To add to the illusion of something happening in the classroom, all characters in the scene would look in the direction of the distraction when it occurred. When the distraction ended, they would return to their slow-motion animations in a seamless manner.

#### b. EEG setup for EEG-NFB sessions

The EEG was acquired from 32 sintered Ag/AgCl multi-electrodes mounted in an elastic cap adapted to children’s head size according to the 10-20 montage (EasyCap GmbH, Germany). One of the electrodes was dedicated to ECG recording and was positioned on the participant’s back on the left under the shoulder plate. Electrodes were equipped with 10 kOhm resistors (20 kOhm for ECG) to avoid induced currents during simultaneous fMRI recordings (Lemieux et al., 1997). To ensure a good electrical contact between the scalp and the electrodes, high-chloride abrasive gel was used after degreasing the scalp with isopropyl alcohol. Before starting the recording, the scalp impedances were lower than 30 kOhm. The EEG was acquired at 500 Hz using a 32-channel BrainAmp MR Plus amplifier (Brain Products GmbH, Germany, RRID:SCR_009443) connected to the cap with a bundled ribbon cable.

The real-time processing of the EEG has been done using OpenVibe (Acquisition Server & Designer 2.2.0, used to acquire EEG and process data respectively, RRID:SCR_014156), an open-source software for Brain Computer Interfaces developed by INRIA (http://openvibe.inria.fr). The following computations were executed on a time moving Hanning window of 3 seconds which was updated every second. First, an average reference was computed by averaging the signal of every EEG electrode and subtracting this average from each electrode. Then, a Fast Fourier Transform was performed on electrode Fz and its spectral power was computed in frequency bands corresponding to theta (4 to 7.5 Hz) and beta oscillations (13 to 19 Hz). Finally, the TBR was computed and the results were sent to the Unity application with the use of a TCP/IP connection. Muscular artefacts due to jaw movements which may corrupt the beta oscillation power were detected by computing the beta oscillation power over the temporal electrode T7. If the beta power of electrode T7 exceeded a threshold of 3 times the beta power of electrode Fz, then the TBR sent to the Unity classroom application was set to zero and the subject was prompted to relax its facial muscles.

#### c. Simultaneous fMRI-EEG setup

Simultaneous EEG-fMRI acquisitions were performed in a 3T MRI (Siemens Magnetom Trio or Siemens Magnetom Prisma Fit). For the EEG, the same setup as for the EEG recordings in a virtual reality environment outside MRI was used but the sampling rate was set to 5 kHZ and a synchronization box was used to assure a 10 MHz alignment to the MRI scanner clock. These changes in EEG recording were necessary for the correction of the artefacts caused by the MRI on the EEG signal. The EEG cap was connected with a short bundle ribbon cable to the amplifier, located at the back of the MRI bore (Jorge et al., 2015). The EEG amplifier and the wires were immobilized using sandbags and memory foam pillow to avoid any vibrations. The child’s head was also restrained with memory foam cushions to limit movement.

A T_2_*-weighted single-shot gradient-echo echo-planar images sequence was used for functional MRI acquisition (TR = 1760 ms, TE = 30 ms, flip angle = 90°, 32 interleaved axial slices of 3 mm with an inter-slice gap of 0.8 mm, 64×64 acquisition matrix, in-plane resolution = 3×3 mm^2^, bandwidth = 2004 Hz/Px, GRAPPA acceleration factor of 2). A 3D T_1_-weighted structural image with a 1 mm isotropic resolution was also acquired.

### 4. Behavioral tasks

#### a. Neurofeedback training task

##### i. Calibration phase

Because the raw TBR of each child is unique, it needs to be normalized (ranging between 0 and 1) in order to adjust distractions and neurofeedback tasks based on each child’s attentional capabilities. A calibration phase has been designed to identify the optimal minimum and maximum values for normalization. This phase consisted of four steps: two relaxing and two focusing periods interleaved, each lasting 20 seconds. For the relaxing periods, the child was prompted by the experimenter to relax, while he/she was prompt to make mental calculation for the focusing periods. During these steps, raw TBR values were collected and sorted, and any values generated by muscle artefacts were removed (see method section §3.b.). The minimum value was calculated as the minimum value of the sorted array, and the maximum value was calculated as the average value of the last quarter of the sorted array.

##### ii. Helicopter display

During each EEG-NFB training, children practiced two runs of EEG-NFB training, with each run lasting for 3 minutes. Those runs were interleaved with cognitive tasks. EEG data were transmitted using TCP/IP protocol to OpenVibe (RRID:SCR_014156) to estimate in real-time the TBR (see method section §3.b.). During these runs, a helicopter was displayed on the white-board of the classroom. The height of the helicopter flight was dependent on the TBR recorded in real-time. The child was asked to find strategies to regulate his/her attention by trying to take off the helicopter to the highest elevation possible and maintain fly at the highest elevation possible.

##### iii. Calculation task as baseline

In this task, children were engaged in solving simple arithmetic calculations. To accommodate the wide range of competencies among children aged 6 to 11 years, we monitored dynamically the performance of the child. The accuracy was computed continuously during the task. When a child achieves a success rate of less than 70%, the difficulty level of the calculations remains the same. Only addition and subtraction operations involving a single digit are presented (e.g., 9 + 4 or 7 - 3). On the other hand, if a subject surpasses a 70% success rate, the task becomes more challenging. Apart from addition and subtraction with single digits, the calculations now include addition and subtraction with two digits, as well as multiplication with one digit (e.g., 34 + 8 or 5 * 9). During each trial, the calculation is presented to the child, who is then required to select the correct answer from three given possibilities. This interactive format actively engages participants and encourages them to focus their attention on the arithmetic. Therefore, we utilized the initial 20 seconds of this cognitively engaging task to establish the TBR baseline (see method section, §6.b.).

#### b. Continuous Performance Task

The Continuous Performance Task (CPT) is a commonly used task for the diagnosis of attention disorders. Children were asked to press a button whenever a letter appeared, except for the letter "X". In each task block, the rate of occurrence of "X" was 25%. A block comprised 20 trials, and each trial lasted 2 seconds (randomized values around: 750ms for fixation cross, 250ms for letter stimuli, and 1000ms for response time). In total, there were six blocks of each condition performed in interplay. If the child did not respond to a correct letter or took more than one second after the presentation of the stimuli to respond it was considered as an omission error, and if the child pressed the button after the letter ’X’ it was considered as a commission error.

#### c. Neurofeedback Transfer task

The NFB transfer task serves as the final assessment in the protocol, evaluating the children’s ability to self-regulate their brain activity without the presence of feedback. During this task, children were instructed to apply the attention regulation strategies learned during EEG-NFB training, but without receiving any visual feedback. Periods of regulation were interleaved with periods of rest every 30 seconds, comprising a total of 6 blocks per condition.

#### d. Behavioral data analysis

##### i. Satisfaction survey

We assessed children satisfaction via a binary response questionnaire given at the end of the protocol. Results are reported in Table 3.

##### ii. CPT

To evaluate the children’s motivation throughout the sessions, their performance in the CPT was measured in terms of accuracy and reaction times. The accuracy was analyzed using a repeated-measure non-parametric ANOVA test (’jmv’ package in R software), to determine the effects of ‘sessions’ and ‘distractor periods’. Reaction time data were cleaned by excluding error trials (mean 10.8% ± 1.7) and trials with reaction times faster or slower than three times the standard deviation from the mean RTs of each individual subject (mean 2.9% ± 0.8). Linear Mixed Model (LMM) approach was employed (’lme4’ package, Bates et al., 2014)) with subject included as a random intercept to evaluate the effects of ‘sessions’ and ‘distractor periods’ as fixed factors. The analysis of variance was performed using the Satterthwaite’s approximation method of degrees of freedom.

### 5. EEG and fMRI data preprocessing

Three children underwent EEG-fMRI scans before and after EEG-NFB training sessions in the virtual reality cave (sessions 2 and 11, respectively). In this preliminary study, only resting-state runs were used to assess data quality, and the transfer run was used to investigate the impact of training on brain activation.

#### a. EEG data in MRI

Gradient artefacts were corrected using a hybrid mean and median average subtraction method, which prevents head movement artefacts from entering the template (Grouiller et al., 2016). For each time point in the template, the values of the *L* neighbouring artifacts were sorted, and the *K* minimal and maximal values excluded from the averaging process (*L* = 30, *K* = 6). This method is less sensitive to motion and is well-adapted to children’s datasets, as it eliminates large outliers from the moving averaged gradient artefact template. Then, the EEG was low-pass filtered with a Finite Impulse Response (FIR) filter with a cut-off frequency of 70 Hz and downsampled to 500 Hz. After detecting the QRS complex in the ECG channel, pulse artefacts were removed using a hybrid mean and median moving average subtraction method *(L = 40, K = 5)*.

Additional preprocessing steps were done with MNE-python (RRID:SCR_005972, Gramfort, 2013) and were similar to those described in the next method section §5.b.

#### b. EEG data in VR

EEG recordings were preprocessed offline using MNE-python (RRID:SCR_005972, Gramfort, 2013). First, the recordings were bandpass filtered, restricting the frequency range to 1-40Hz using a FIR filter. A notch filter was also applied to eliminate 50Hz line noise. Second, to ensure data quality, segments containing artifacts and channels with poor signal were manually annotated and excluded from subsequent analyses. To remove noncerebral artifacts such as eye blinks and muscle activity, Independent Component Analysis (ICA) decomposition was performed using the Picard algorithm (Ablin et al., 2018). Finally, bad channels were interpolated using spherical spline interpolation. Recordings were re-referenced to the average reference for further analysis.

#### c. fMRI data

Both structural and functional MRI data were preprocessed using *fMRIPrep* 21.0.2 (Esteban et al., 2019) on *Nipype* 1.6.1 (Gorgolewski et al., 2011).

##### i. Anatomical data

T1-weighted (T1w) structural images were corrected for intensity non-uniformity (*N4BiasFieldCorrection, ANTs 2.3.3,* Avants et al., 2008; Tustison et al., 2010), and skull-stripped (*antsBrainExtraction.sh, ANTs 2.3.3*) using OASIS30ANTs as target template. Brain tissue was segmented into cerebrospinal fluid (CSF), white-matter (WM) and gray-matter (GM) (*fast, FSL 6.0.5.*, Zhang et al., 2001). Volume-based spatial normalization to standard spaces (MNIPediatricAsym:cohort-3, an *unbiased template for pediatric data from the 4.5 to 18.5y age range*) was performed through nonlinear registration (*antsRegistration, ANTs 2.3.3*), using brain-extracted versions of both T1w reference and the T1w template.

##### ii. Functional data

For each subject, the functional MRI (i.e. the two resting-state runs and the NFB transfer run)underwent the following pre-processing steps. First, a reference volume and its skull-stripped version were generated using a custom methodology of fMRIPrep. Rigid-body head-motion parameters with respect to the BOLD EPI reference were estimated (*mcflirt, FSL 6.0.5.*, Jenkinson et al., 2002) before any spatiotemporal filtering. BOLD runs were slice-time corrected (*3dTshift, AFNI*R, RID:SCR_005927, Cox, 1996). The BOLD time-series were resampled onto their original, native space by applying the transformations to correct for head-motion. The BOLD reference was then co-registered to the T1w (*mri_coreg, FreeSurfer*; *flirt, FSL*) with the boundary-based registration cost-function. Co-registration was done with six degrees of freedom.

The data underwent additional preprocessing using the AFNI software (RID:SCR_005927, Cox, 1996). A spatial smoothing was applied using 8-mm FWHM Gaussian kernel to reduce surrounding noise (Op De Beeck, 2010). Finally, a linear regression was performed to eliminate nuisance variables such as estimated motion parameters, the first-order temporal derivatives of motion parameters, squared motion parameters, and mean time courses of the cerebrospinal fluid and white matter signals. These nuisance variables were further adjusted for linear and higher order polynomial trends (Power et al., 2012; Van Dijk et al., 2012).

### 6. EEG and fMRI data analysis

#### a. Resting-state fMRI networks

To identify brain networks, we applied a probabilistic independent component analysis (ICA) approach using the FMRIB Software Library’s MELODIC toolbox on the full dataset, obtained by temporally concatenating across subjects, sessions and runs (Schmithorst and Holland, 2004) to create a single four-dimensional dataset decomposed into 20 independent components. We categorized the resulting ICs into three groups (akin to Hadj-Bouziane et al., 2014; Guedj et al., 2016) based on visual inspection: 1) ‘real’ (bilateral networks consistent with known anatomical and functional circuits), 2) ‘duplicate/unclear’ (bilateral or noisy map hat overlapped with real networks), and 3) ‘noise-related’ ICs (random speckle patterns and rim patterns). To produce final maps, we applied a posterior probability threshold of p < 0.5 (equal loss placed on false positives and false negatives) by fitting a Gaussian/Gamma mixture model to the histogram of intensity values (Beckmann and Smith, 2004). Similarly, to evaluate data quality at the individual level, we also performed ICA for each subject separately concatenating pre-and post-resting-state fMRI runs.

#### a. EEG Microstates

The EEG microstates analysis was conducted using Pycrostates (Férat et al., 2022a). For each recording, the local maxima of the global field power (GFP), which represent segments of the EEG data with the highest signal-to-noise ratio (Koenig and Brandeis, 2016), were identified. These GFPs were then subjected to modified k-means clustering (polarity-independent) with 100 repetitions, selecting the clustering result with the highest global explained variance. A predetermined value of k = 5 microstate maps (cluster centroids) was estimated for each recording, considering the exploratory nature of the study and drawing from previous literature (Michel and Koenig, 2018; Férat et al., 2022b).

The estimated maps at the recording level were subsequently merged and subjected to the same clustering algorithm to extract k = 5 topographies that best represent the dataset. These k = 5 global dominant topographies were then fitted back to the original EEG data. In this procedure, time points were assigned cluster labels (i.e., microstate topographies) based on spatial correlation analysis. Each time point was assigned to the topography with which it had the highest absolute spatial correlation. To ensure temporal continuity, a smoothing window of 50ms was applied, adjusting the central time point’s correlation with a smoothing factor of 10.

#### b. TBR in EEG-NFB sessions

Frequency bands were obtained by applying spectral decomposition techniques using Fast Fourier Transform algorithm (Welch, 1967). Specifically, we focused on the theta and beta band to further compute TBR. The average TBR on Fz electrode was calculated across all eight sessions for each subject, for the two helicopter task runs and the first 20 seconds of the calculation task, which served as the baseline. Furthermore, we calculated the evolution of the TBR relative to baseline for each session of every subject as followed:

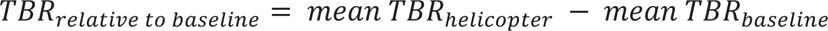

Finally, the TBR evolution was averaged across all subjects, enabling us to gain insights into the collective trends and variations in TBR across the entire child cohort.

#### c. Transfer Run

##### i. EEG

The data were divided into 10-second epochs and categorized as either ’Rest’ or ’Regulation’, depending on the corresponding block period of the NFB transfer task. Epochs containing residual artifacts were manually identified and excluded from further analysis. Similar to the TBR computation during EEG-NFB sessions, we used Fast Fourier Transform algorithm to extract the theta and beta frequency bands. Subsequently, we calculated the average TBR on the Fz electrode separately for the ’Rest’ and ’Regulation’ epochs.

##### ii. fMRI

The linear regression model described above (see section method §5.1.ii) included an additional regressor representing the onset of ’Rest’ and ’Regulation’ blocks. Contrasts comparing ’Regulation’ to ’Rest’ conditions were computed for each child and subsequently entered into a second-level analysis using a one-sample Analysis Of Variance (ANOVA). A statistical threshold of p < 0.05, uncorrected for multiple comparisons, was applied in the analysis.

## RESULTS

### 1. Satisfaction and sustaining motivation

At the end of the experiment, a satisfaction survey was administered to the six participating children. The results, displayed in Table 3, indicate that all children reported enjoyment of the study. Furthermore, 5 out of the 6 children expressed satisfaction with the virtual reality experience and noted improvement in their concentration abilities. However, it was found that 83% of the children were unable to consistently apply the concentration strategies learned during the sessions in their daily classroom setting.

To assess the maintenance of the children’s motivation during the EEG-NFB sessions in the VR, we evaluated their performance in the CPT throughout the sessions in terms of accuracy and reaction times. The integration of distractions was a critical aspect of the experimental design, as it mimicked the conditions of a real-life classroom and challenges the participants’ attentional capacities. As such, we also studied the effects of these distractions on the children’s performance. The results of the study showed that the four participating children had an average accuracy of 89% (±5%) on the CPT task across the eight runs (Figure 3A). There was no significant impact of session or distractor period on accuracy (**χ**2(15)=20.79; p=0.14). However, a main effect of session (**χ**2(7)=29.80; p<0.0001) and distractor period (**χ**2(1)=19.74; p<0.0001) was observed in the analysis of reaction times (Figure 3B), without any interaction between the two factors. Post-hoc tests revealed that children were slower to respond during distractor periods compared to the period without distractions, and they were significantly slower during the session 4 compared to session 1 and session 3.

**Figure 3.**
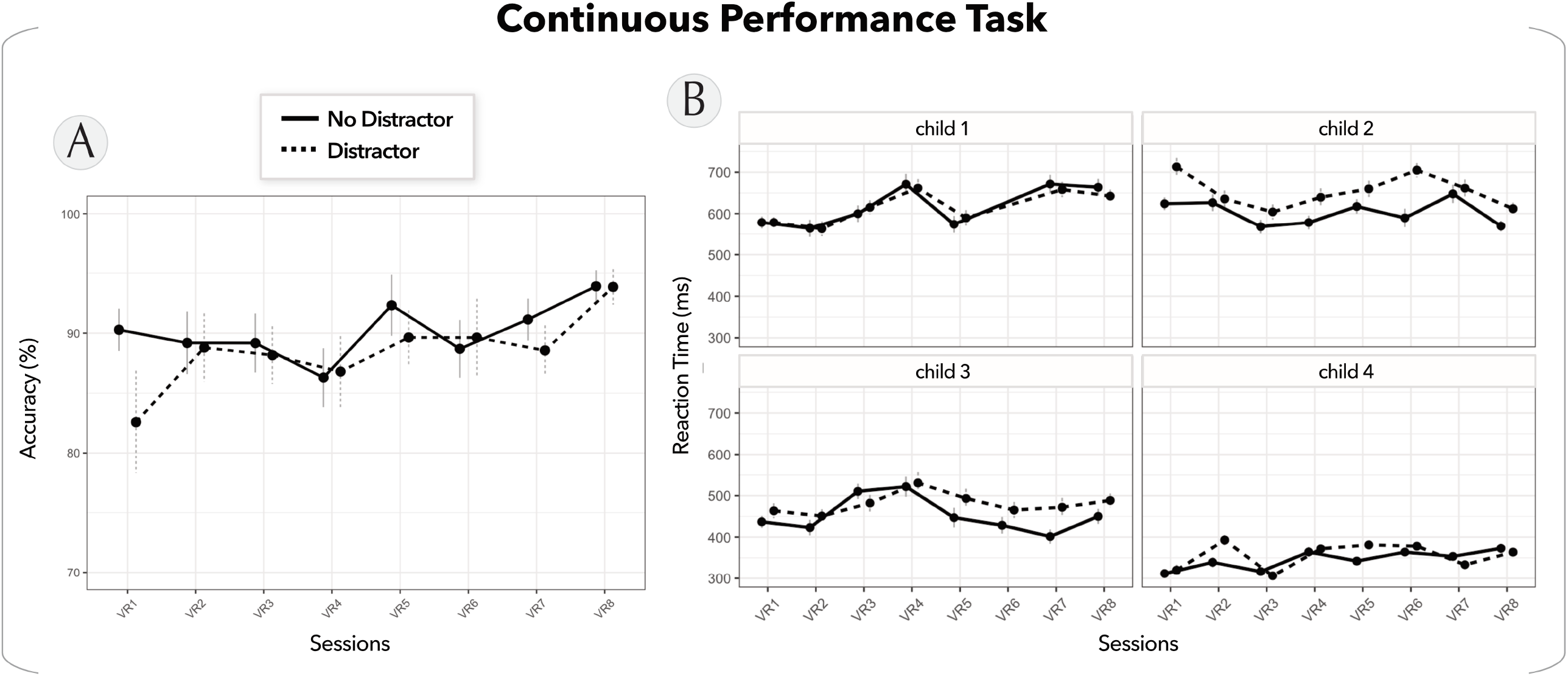

Taken together these results demonstrate the stability of the children’s performance across the multiple sessions, indicating a sustained level of motivation to complete the experimental protocol. Additionally, the findings highlight the impact of the distraction period, confirming the expected distraction effect.

### 2. Feasibility in children: quality assessment of simultaneous EEG-fMRI data

To ensure the feasibility of our protocol, we examined the quality of the EEG and fMRI data acquired during simultaneous recordings (sessions 2 and 11). High-quality data is essential for future studies to evaluate the effect of neurofeedback on brain activation and connectivity.

To assess data quality, we focused on resting-state runs. First, we applied the ICA approach to fMRI data to identify the well-known and documented resting-state networks (Smith et al., 2009). At the group level, we observed seven resting-state networks that exhibited the typical bilateral pattern of ’real’ resting-state networks (Figure 4A). We categorized the remaining networks as either ’duplicate/unclear’ or ’noise-related’ ICs (Supplementary Figure S1). At the individual level, the ICs were less clear, likely due to the smaller amount of data and participant movements during the scanning session (680 time points versus 2040 for the grouped data). However, we were still able to identify the dorsal fronto-parietal network involved in attention processes (Corbetta and Shulman, 2002) at both the group and individual level (Figure 4B).

**Figure 4.**
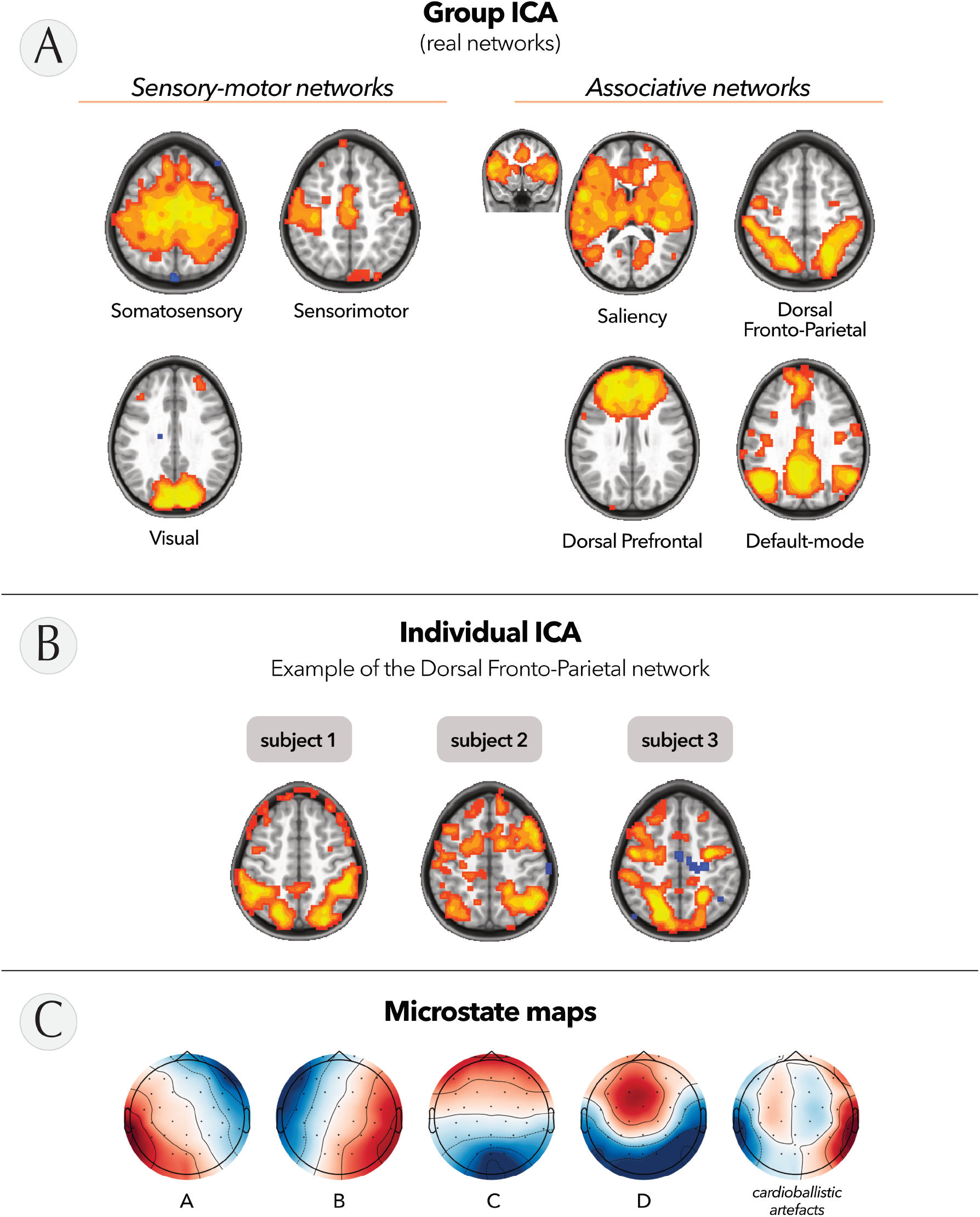

In addition, we evaluated the quality of the EEG data acquired simultaneously with fMRI by performing microstate analysis at the group level. Similar to ICA, microstate analysis is a technique that enables the examination of spatiotemporal patterns within EEG recordings through k-means clustering. This method involves decomposing the multichannel EEG signal into a series of quasi-stable states, where each state is characterized by a specific spatial distribution of scalp potentials known as a microstate map. This analysis provides valuable insights into the temporal dynamics and spatial patterns of the EEG data and can be used in a future study to evaluate the impact of EEG-NFB training on these dynamics. As depicted in Figure 4C, decomposition of the signal resulted in five maps. Four of these maps were consistent with the microstates traditionally described in the literature (Michel and Koenig, 2018), while the fifth topography corresponded to residual cardioballistic artifacts originating from the simultaneous EEG-fMRI setup (Iannotti et al., 2015).

### 3. Potential of EEG-NFB combined with VR: preliminary findings

Finally, we assessed the potential of this innovative protocol. First, we examined the EEG data recorded during the EEG-NFB training sessions. The average TBR on Fz electrode was computed per session for each child, relative to their baseline. The progression of this value across the 8 training sessions for the four children is depicted in Figure 5 (left panel). No significant effect was observed when considering the session factor. However, a notable trend emerges when comparing the first training period (session 1 to 4) to the second (session 4 to 8). Three out of four children demonstrated a decrease in their TBR during the later stage of training, indicating a learning process for controlling brain activity (see Figure 5, right panel).

**Figure 5.**
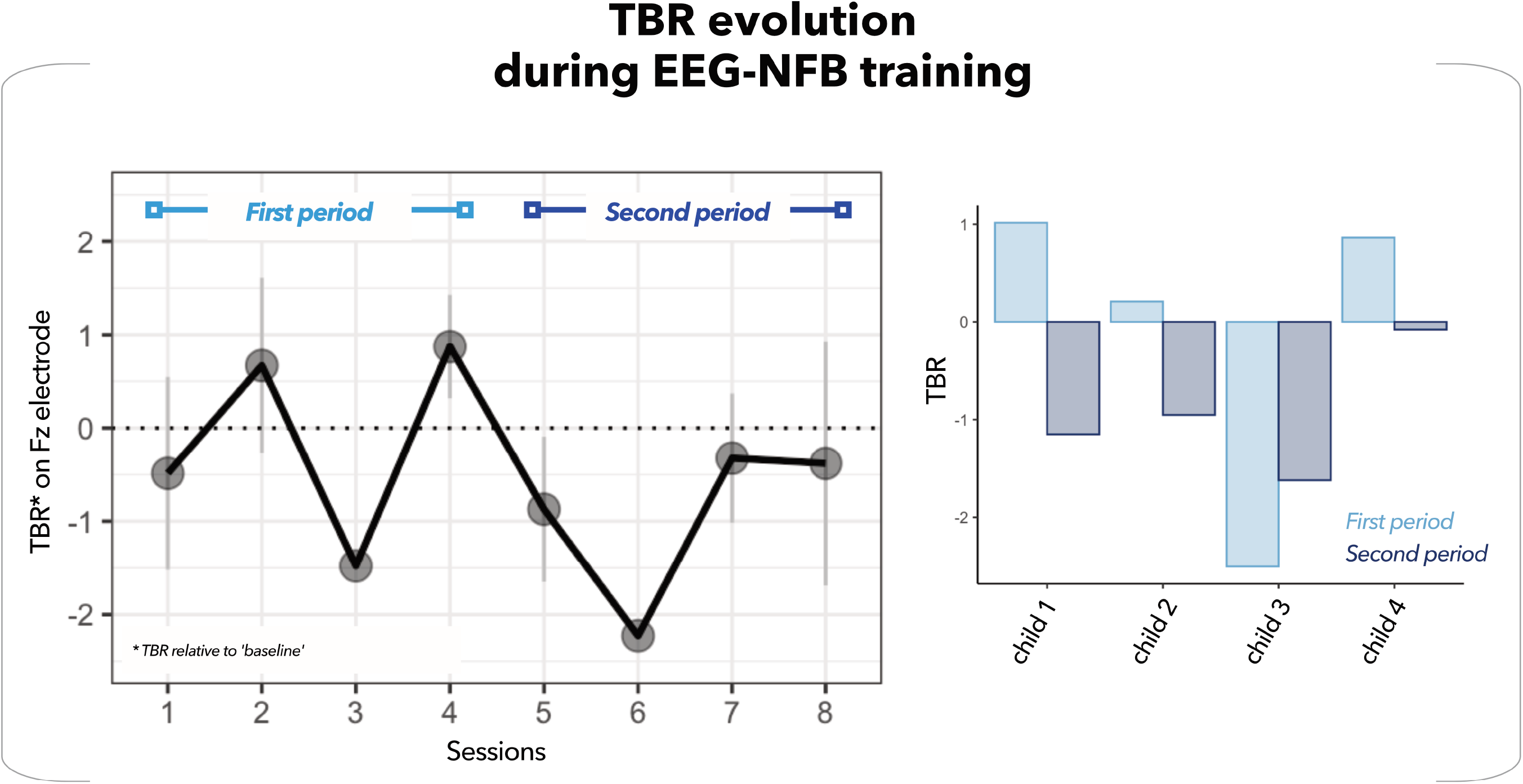

Second, we measured the children’s ability to regulate their attention during the NFB transfer task, i.e. inside the MRI scanner and without any feedback. Our objectives were to evaluate their proficiency in controlling brain activity and to investigate the corresponding neural correlates. To do so, the children were instructed to alternate between periods of rest and periods of concentration. In the EEG data, we observed a consistent (not significant) decrease in TBR at the Fz electrodes for all three children (Figure 6A). Interestingly, we observed that the children did not exhibit a uniform pattern of TBR modulation. Child 1 displayed a decrease in theta activity, while child 2 displayed an increase in beta activity, and child 4 showcased the ability to modulate both theta and beta rhythms.

**Figure 6.**
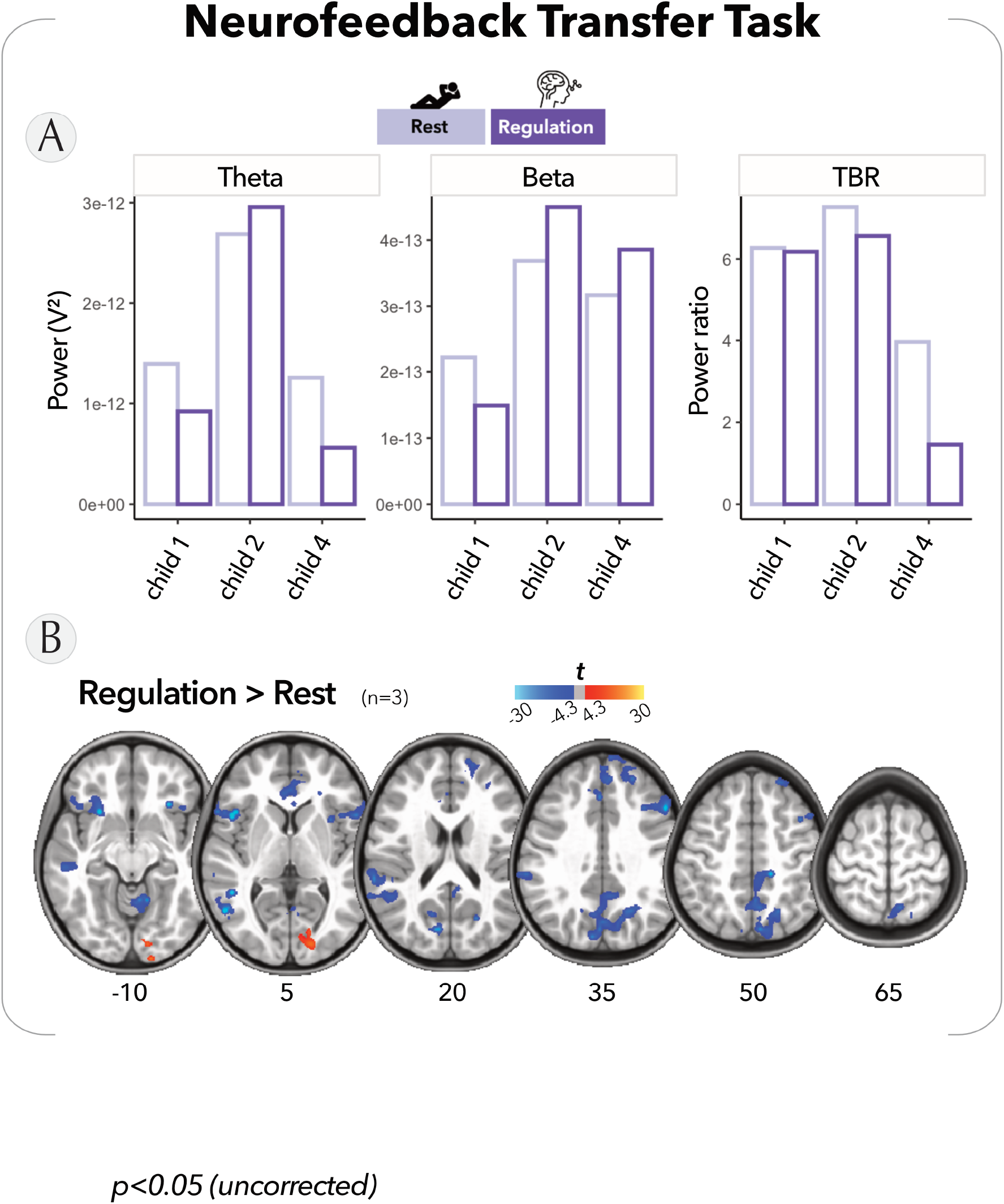

Simultaneously, in the fMRI data, we found a significant reduction in brain activation within regions associated with the default-mode network and sensorimotor cortices, during the attention regulation periods compared to the resting periods (although not corrected for multiple comparisons) (see Figure 6B and Supplementary Material Table S1). This finding indicates a cognitive engagement in the children and potentially suggests an improved ability to filter out external distractions.

## DISCUSSION

The answer to our initial question is yes, our innovative EEG-NFB protocol, combined with a virtual reality environment, is feasible in children with ADHD and is likely to be accepted and attractive to this vulnerable population. Additionally, to the best of our knowledge, this is the first study to combine these technologies to propose a non-pharmacological alternative for treating attention deficits. This report also provides comprehensive information about the VR environment developed by our team, specifically designed to introduce EEG-NFB training to school-aged children between 6 and 11 years old.

Among the children who began the EEG-NFB training sessions (n = 4), the final participation rate in the study was 100%. One family attended the initial neuropsychological assessment session but ultimately declined to participate, due to time constraints. These findings demonstrate that once enrolled in the study, and particularly once the EEG-NFB training commenced, participants and their families exhibited a strong interest and willingness to fully engage in the intervention. While comparisons with other studies may be challenging due to the small cohort size utilized in this study, the compliance results appear to surpass those reported in other NFB studies involving children (e.g. Gevensleben et al., 2010).

The satisfaction results of this study are promising, but it is important to consider the limitations within its context. Firstly, our satisfaction scale was limited to multiple choice responses, which could benefit from the use of a numerical scale for a more precise quantification of children’s satisfaction. Secondly, the satisfaction survey indicated that the children encountered difficulties in applying the concentration strategies they had learned to control the helicopter in a real classroom environment. In future research, we intend to enhance the neurofeedback intervention by incorporating a pre-session tutorial. This tutorial will provide children with a deeper understanding of attention processes and guide them on effectively utilizing these strategies to achieve their goals. By addressing these limitations, we aim to further optimize the intervention’s efficacy and improve overall satisfaction among the participants.

The quality of both the MRI and EEG recordings was deemed adequate for conducting the intended group analyses and evaluate the effects of our innovative EEG-NFB protocol. However, at the individual level, the analysis was somewhat constrained by residual movement artifacts. Nonetheless, in future studies, enhancing data quality, particularly during resting runs (i.e. when no active engagement of the participant is required), could be achieved by using visual stimulation with abstract shapes. It has been shown that this approach can improve children’s compliance during prolonged sessions, such as resting-state fMRI, thereby improving data quality (Vanderwal et al., 2015).

To conclude on the therapeutic potential of our protocol, it is essential to acknowledge the limitations of our study, namely the small sample size of our group of children and the absence of attentional or neuropsychological disorders among the four participants. Despite these limitations, it is encouraging to observe that both EEG and fMRI data consistently align in the same direction, indicating an improvement in TBR. The reproducibility of this improvement even in the absence of feedback highlights the children’s capacity to acquire the skill of controlling their own brain activity. Furthermore, this is also confirmed by neuroimaging data showing a decrease in activation in regions of the default-mode network, which operates antagonistically to the task-positive network, also referred to as the attention network (Greicius et al., 2003; Fox et al., 2005).

An important aspect of this study involves the individual variations in TBR modulation observed during the transfer run. This variability underscores the well-known complexity inherent in the treatment of children with ADHD. The syndrome often manifests as multidimensional, with significant heterogeneity in symptoms and electrophysiological profiles (Clarke et al., 2001; Luo et al., 2019). These findings reinforce the notion that personalized interventions, potentially based on individualized electrophysiological markers, are crucial for effective therapy of this pathology.

## Conclusion

To the best of our knowledge, this study represents a pioneering effort in combining EEG-NFB and VR to provide a relatively short and playful therapeutic protocol aimed at teaching children with ADHD to regulate their own brain activity.

Once enrolled, the full rate of engagement in the intervention program also shows that the children largely accepted and enjoyed the program. While the efficacy of the protocol has yet to be validated, the preliminary findings are exceptionally promising.

To summarize, this novel EEG-NFB protocol combined with VR holds significant potential as a valuable tool for enhancing executive functions and behavioral skills among children with ADHD. Considering the power of brain plasticity (Loriette et al., 2021), this innovative intervention could have a significant impact on the well-being and development of people with ADHD. In perspective, this study not only highlights the initial success of the protocol, but also emphasizes the need to explore playful and immersive interventions in the treatment of childhood pathologies.

## ACKNOWLEDGEMENTS

This work was supported by a FreeNovation grant from Novartis Research Foundation, by a grant from Swiss National Science Foundation (SNSF grant 188769 to F.G.), and by a grant from the Gertrude Von Meissner Fondation (to C.G.). This study was conducted using the imaging platform and virtual reality platform of the Brain and Behaviour Laboratory (BBL), Cognitive and Affective Neuroimaging Section of the Center for Biomedical Imaging (CIBM MRI UNIGE).

## Notes

### Competing Interest Statement

The authors have declared no competing interest.

