## Supplementary_material for "Self-regulation of attention in children in a virtual classroom environment: a feasibility study"

Figure S1. **Example of other ICs**

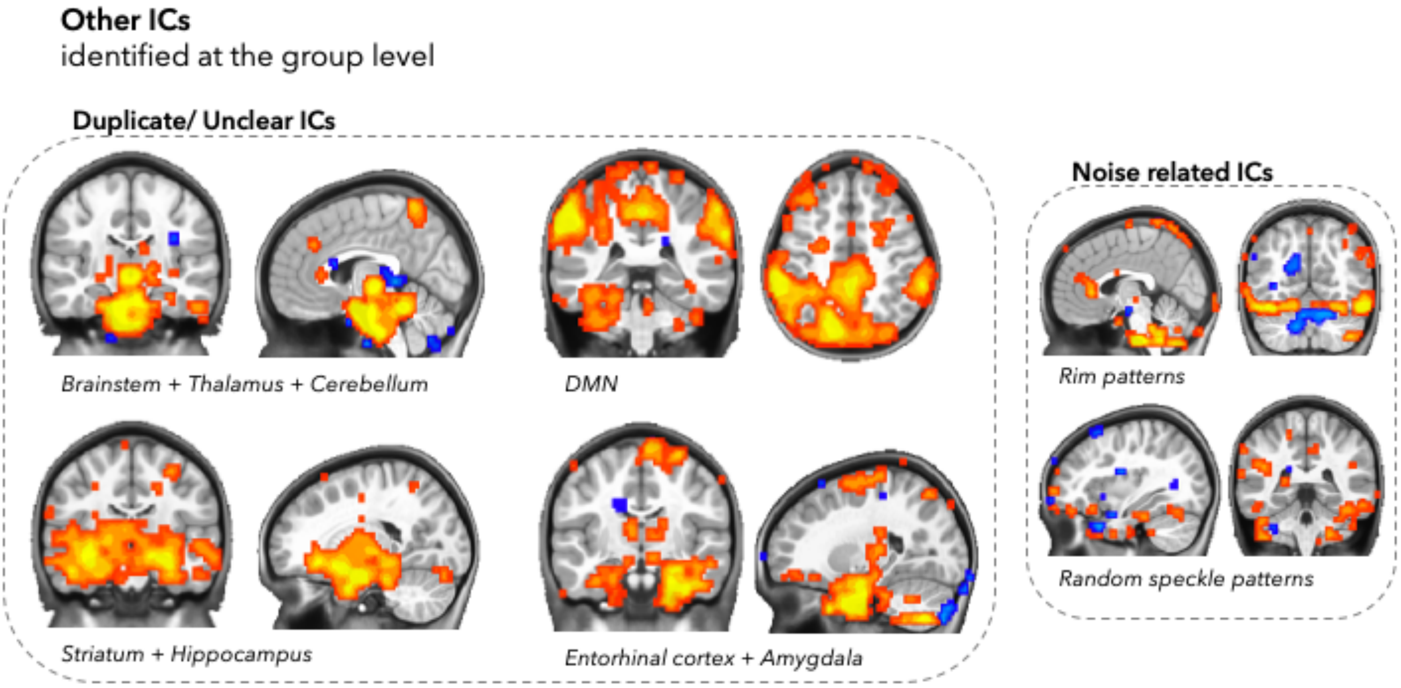


*Table S1.* ***Cluster Table***

| Cluster size | x | y | z | Regions |
|---|---|---|---|---|
| 623 | 9.5 | -83.5 | 55 | R Precuneus |
| 310 | -8.5 | 45.5 | 13.8 | L Superior Frontal Gyrus |
| 234 | -68.5 | -53.5 | 6.2 | L Middle Temporal Gyrus |
| 153 | -50.5 | 30.5 | -8.8 | L Inferior Frontal Gyrus |
| 101 | 69.5 | -32.5 | 32.5 | R SupraMarginal Gyrus |
| 92 | 60.5 | 27.5 | 2.5 | R Inferior Frontal Gyrus |
| 64 | 9.5 | -86.5 | 2.5 | R Calcarine Gyrus |
| 53 | 54.5 | 27.5 | 36.2 | R Middle Frontal Gyrus |

x, y, z coordinates refer to the peak location in MNI space (LPI orientation). The cluster size refers to the number of voxels. Extent threshold of 50 voxels (face+edge+cornerwise neighbors). L, left; R, right.
