## Supplementary material for "Self-regulation of attention in children in a virtual classroom environment: a feasibility study": Legends_figures_tables

### FIGURE LEGENDE

*Figure 1.* ***Protocol overview***.

*Figure 2.* ***Virtual classroom environment***. (A) View from the front. (B) View from the back. (C) Child's installation within the VR cave. An EEG cap is placed on the child's head, complemented by a headband equipped with infrared sensors. These sensors enable the visual scene to dynamically synchronize with the child's movements, enhancing the overall realism of the VR experience.

*Figure 3.* ***Continuous performance task***. (A) Task accuracy average across the four children and (B) Reaction time for each children, during sessions in the VR classroom environment. For the condition without distraction (solid lines) and with distractions (dotted lines). Error bars reflect the standard errors of the mean.

*Figure 4.* ***fMRI and EEG data quality***. (A) Resting-state networks identified in fMRI with ICA at the group-level and overlaid on axial or coronal slices. Warm colors correspond to positive correlations and cool colors to negative correlations. (B) Associated dorsal fronto-parietal network from the subject-level. (C) Microstate maps identified in EEG acquired simultaneously with fMRI at the group-level.

*Figure 5.* ***TBR evolution during EEG-NFB training***. [Left panel] Average TBR on Fz electrode for the eight EEG-NFB training sessions. This TBR value corresponds to the TBR helicopter task value after subtracting the baseline. [Right panel] TBR values extracted for each child from the first (light blue) and second (dark blue) period of the protocol.

*Figure 6.* ***EEG data quality***. (A) Theta and Beta power, and TBR on Fz electrode extracted for each child from the 'Rest' (light purple) and 'Regulation' (dark purple) periods. (B) Statistical maps of the contrast 'Regulation vs. Rest' overlaid on axial slices. The color-bar shows T-values task-related activations (yellow/red) and inactivation (blue).

### TABLES

*Table 1.* ***Unity Assets used in the VR simulation***

| **Name** | **Description** | **Asset link** |
|---|---|---|
| **School classroom** | Classroom used as the 3D environment of the VR simulation | *https://assetstore.unity.com/packages/3d/characters/humanoids/2-toon-people-116917* |
| **Toon characters** | 20 'toon kids' (10 boys and 10 girls) sat at their desks in pairs, all around the (participant) child. The latter was placed in the center of the virtual classroom sitting at a real school desk (which was the center of the CAVE system). | *https://assetstore.unity.com/packages/3d/characters/humanoids/humans/toon-kids-55945* |
| **Toon People** | 2 'toon peoples' were used to represent the virtual mistress and school's headmaster. | *https://assetstore.unity.com/packages/3d/characters/humanoids/2-toon-people-116917* |
| **Everyday Motion Pack Free** | Package of animations (idle, sit, walk, talk) used to animate virtual character bodies. | *https://assetstore.unity.com/packages/3d/animations/everyday-motion-pack-free-115067* |

| | | |
|---|---|---|
| **SALSA Lip Sync** | Plugin used to animate virtual character faces, to generate various random head and gaze directions, eye-blinks, “look at target” behaviours. It also allowed virtual characters’ lips moving in synchrony with virtual character’ speeches (audio-to-speech). | *https://assetstore.unity.com/packages/tools/animation/salsa-lipsync-suite-148442* |

*Table 2.* ***Panel of distractions***

| Character | Type of stimulation | Description |
|---|---|---|
| Mistress | Audiovisual | - walk,<br>- cough<br>- yawn,<br>- speak randomly among 11 sentences (voices recorded from one adult female) |
| Headmaster | Audiovisual | - enter and exit the classroom,<br>- walk,<br>- cough,<br>-yawn,<br>- speak randomly among 6 sentences (voices recorded from one adult male) |
| Kids | Visual | - raise their hands |
| | Audiovisual | - speak among 53 sentences (voices recorded from one girl and one boy) |
| Other | Audiovisual | - phone on the mistress’ desk is vibrating |
| | Audio | - end of class bell rings |
| | Audiovisual | - noisy insects (fly, dragonfly, butterfly) fly in front of the whiteboard |

*Table 3.* ***Satisfaction survey***

| Questions | Yes | No | Sometimes |
| --- | --- | --- | --- |
| Did you like the VR classroom sessions? | 83% | 0% | 17% |
| Were you happy to come and do the experiment? | 100% | 0% | 0% |
| Do you feel that you have improved your ability to fly the helicopter? | 66% | 0% | 33% |
| Did you have a strategy for flying the helicopter? * | 100% | 0% | 0% |
| Do you use these strategies to focus in class? | 17% | 83% | 0% |
| Do you feel like you are able to focus better? | 83% | 0% | 17% |

*example of strategies used: "I recited the multiplication tables" ; "I focused on the image of the helicopter" ; "I focused on a detail of the classroom".

The shaded cells correspond to the majority response over the 6 children who completed the protocol. Note that 2 of them were removed from the rest of the analyses due to technical issues (see method section, §2).
